# Diversification is correlated with temperature in white and sulfur butterflies

**DOI:** 10.1101/2022.09.22.509088

**Authors:** Ana Paula S. Carvalho, Hannah L. Owens, Ryan A. St Laurent, Chandra Earl, Kelly M. Dexter, Rebeccah L. Messcher, Keith R. Willmott, Kwaku Aduse-Poku, Steve C. Collins, Nicholas T. Homziak, Sugihiko Hoshizaki, Yu-Feng Hsu, Athulya G. Kizhakke, Krushnamegh Kunte, Dino J. Martins, Nicolás O. Mega, Sadaharu Morinaka, Djunijanti Peggie, Helena P. Romanowski, Szabolcs Sáfián, Roger Vila, Houshuai Wang, Michael F. Braby, Marianne Espeland, Jesse W. Breinholt, Naomi E. Pierce, Akito Y. Kawahara, David J. Lohman

## Abstract

Temperature is thought to be a key variable explaining global patterns of species richness. However, to investigate this relationship carefully, it is necessary to study clades with broad geographic ranges that are comprised of species inhabiting diverse biomes with well-characterized species ranges. In the present study, we investigate the link between temperature and diversification in the butterfly family Pieridae (sulfurs and whites) by combining Next Generation sequences and published molecular data with fine-grained distribution information. After building the most comprehensive phylogeny of the group, with almost 600 species and all higher taxa (subfamilies, tribes and subtribes), we found strong support for the following relationships within the family: Dismorphiinae + (Coliadinae + (Pseudopontiinae + Pierinae)). With a curated distribution dataset of over 800,000 occurrences, we conducted multiple comparative phylogenetic analyses that provided strong evidence that species in environments with more stable daily temperatures or with cooler maximum temperatures in the warm seasons have higher diversification rates. We also found a positive correlation between speciation and extinction with paleotemperature: as global temperature decreased through geological time, so did diversification rates. Although many studies demonstrate higher diversity in the tropics, we have been able to identify specific climate variables associated with changes in diversification, while also inferring the most robust and well sampled phylogenomic framework for Pieridae to date.

## Introduction

Understanding factors that affect variation in species diversity across Earth is a central pursuit of ecology (1). While multiple biotic and abiotic factors drive speciation and extinction (2–5), temperature is thought to be a key variable impacting diversity patterns (6–8). Diversity is expected to be greater in warmer environments due to higher evolutionary rates (the evolutionary rate hypothesis (2)). According to this hypothesis, species in warmer areas tend to have higher mutation rates, faster metabolism, and shorter generation times, all of which are directly associated with increased rates of evolution (2, 4, 9–11). Due to the lack of pronounced temperature variation in species-rich areas (*e*.*g*., some tropical regions), authors also hypothesize that temperature stability affects diversification (12). This relationship is based on the premise that climatic stability favors the evolution of specialized adaptations (climatic stability hypothesis (12)). The inherent temperature stability of tropical climates compared to temperate ones is therefore thought to reduce extinction rates.

To test the link between temperature and diversification, it is necessary to study widely distributed clades of species inhabiting diverse biomes that have sufficiently fine-grained locality data. Additionally, a well-supported, dated phylogenetic tree is necessary to account for relationships among species when evaluating patterns of diversity. Although many studies have assessed the correlation between ambient temperature and diversity, these studies have primarily focused on vertebrates (3, 13–19). Considerably less attention has been directed to other, substantially more diverse groups like insects, which–as ectotherms–have life histories more likely to be intrinsically connected to temperature (20). Butterflies are among the best documented insect groups, with nearly 34.5 million locality records on the Global Biodiversity Information Facility (GBIF, as of September 2022).

Studies of butterfly diversity have largely focused on the coevolution with their host plants (21). Prior investigations have found strong evidence for reciprocal adaptations between Pierinae butterflies and their host plants as predicted by an escape-and-radiate coevolutionary scenario (22, 23). However, an extensive study examining evolutionary patterns of host use across all butterflies found weak evidence for coevolution in many clades (24). Host plants are just one factor that might affect insect diversification rates; less attention has been paid to the role of climate.

Butterfly species richness is generally much higher in equatorial regions than temperate regions (25), and is also correlated with ambient temperature (26). Like other ectotherms, temperature strongly influences insect larval development and diapause (27). Ambient temperature and sunlight have direct effects on behaviors such as basking, and indirect effects on life history traits including phenology and voltinism (*e*.*g*., 28, 29). Consequently, extreme or highly variable temperatures can negatively impact fitness (30). Temperature mediated diversification is another important context in which to study diversification dynamics in ectothermic organisms, such as butterflies.

The butterfly family Pieridae comprises 1,159 species in 86 genera distributed across the globe (31). Most of their diversity is concentrated in the tropics (32), but Pieridae is known for having species adapted to extreme cold or dry conditions, such as the Arctic (33) and deserts (34). Pierid monophyly is well established (35, 36), but relationships among subfamilies and tribes remain contentious (see 37–39 for clade or region specific studies) (Fig. 1). A comprehensive phylogeny of Pieridae will not only disentangle poorly understood relationships but also illuminate persistent questions regarding the role of thermal ecology in insect macroevolution. Here we construct a comprehensive species-level dated tree of Pieridae with 593 species from 84 genera and infer the evolution of host plant interactions. We combine our phylogenetic reconstruction with climate data extracted using distribution records to examine whether speciation and extinction are correlated with 1) warmer climates (annual mean temperature), 2) seasonality and daily temperature variability (temperature annual range and annual mean diurnal range), 3) extreme temperatures (maximum temperature of the warmest month and minimum temperature in the coldest month), and/or 4) paleotemperatures.

**Fig. 1.**
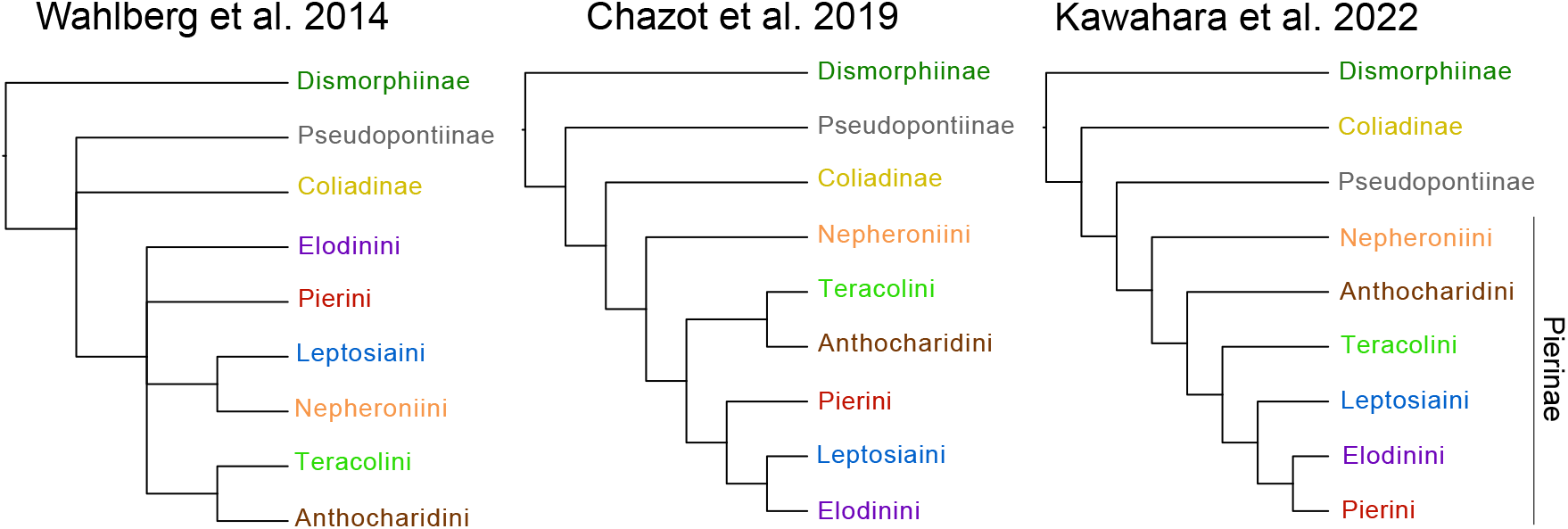
Summary of the branching patterns of Pieridae among recent molecular phylogenetic studies.

## Results

### Diversity patterns and temperature

Pieridae lineages have steadily increased since the family’s origin around 70 Ma ago, according to the deterministic lineage through time plot (dLTT) (Fig. S1). Fitted pulled speciation rates (PSR) gradually decreased from the stem lineage until 40 Ma, when rates started to increase and had two peaks between 20 Ma and the present (Fig. S2).

The BAMM (Bayesian Analysis of Macroevolutionary Mixtures) estimation of net diversification rates identified five clades with increases in diversification rates: *Colias*; a subclade of *Catasticta* (+ *Archonias*); *Delias*; and clades within *Pieris* and *Mylothris* (Fig. 2, Fig. S3). These patterns were consistent with our RevBayes analysis (Fig. S4). The distinct shift configuration of BAMM suggests a high (>0.75) marginal probability of shifts for *Colias, Pareronia*, and a subclade of *Catasticta* (+ *Archonias*) (Fig. S3).

**Fig 2.**
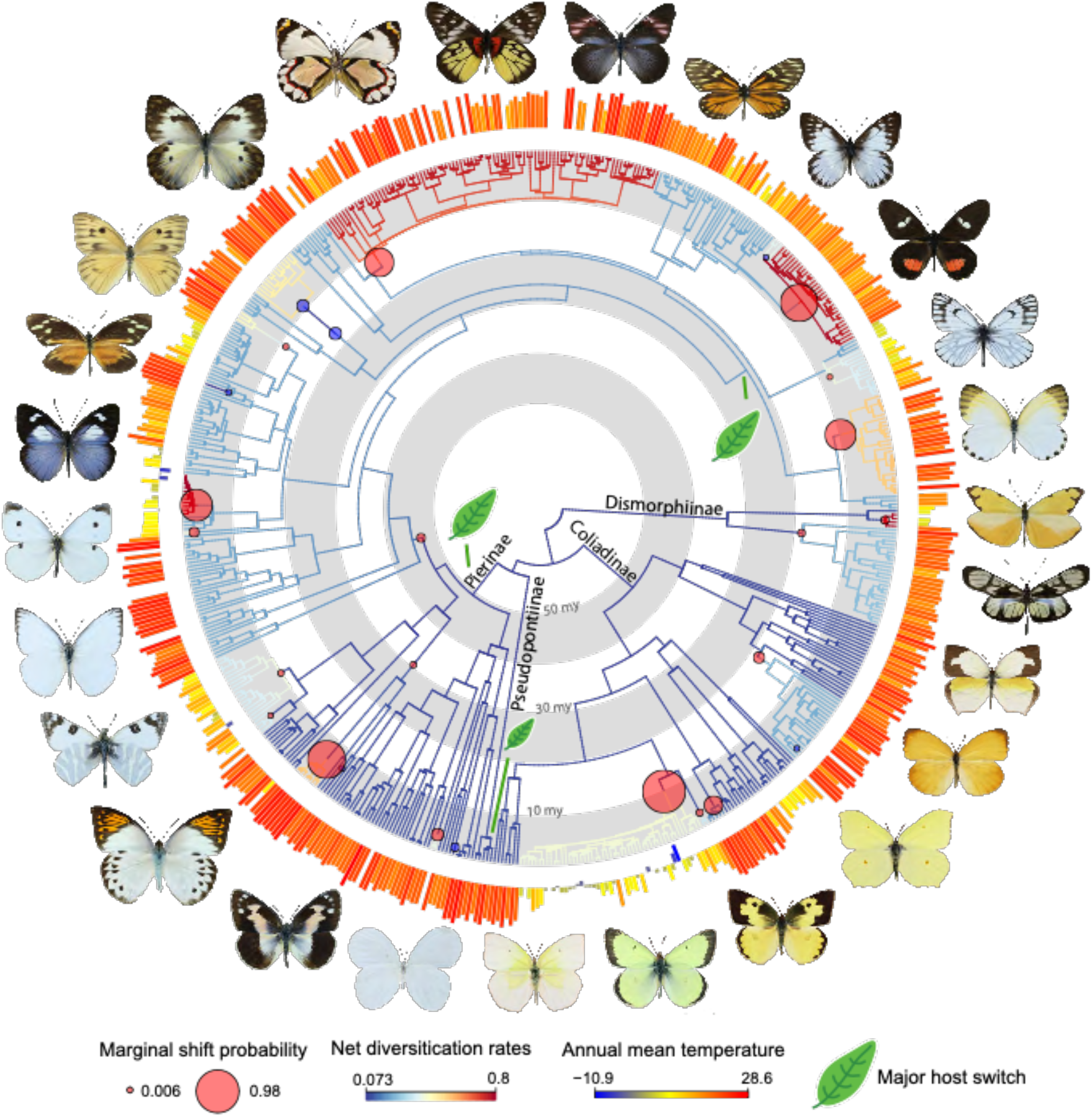
Evolutionary relationships and diversification patterns of Pieridae. Time-calibrated tree of 593 species. Branches with significant net diversification rate shifts as estimated by BAMM are indicated with red and blue circles. Annual mean temperature (BIO1, WorldClim) for each species is indicated with colored bars in the outer ring. A green leaf indicates a major shift in host plant use along the branch as reconstructed by SIMMAP. Butterfly images are not to scale.

Our phylogenetic generalized least squares (PGLS) analyses found significant relationships between temperature and speciation/extinction for two variables: 1) minimum temperature of the coldest month (positive for speciation, negative for extinction), and 2) temperature annual range (negative for speciation, positive for extinction). We also found a significant negative relationship between speciation and mean diurnal range (Fig. S5). The annual mean temperature and maximum temperature of the warmest month did not show significant relationships with either speciation or extinction.

However, when QuaSSE (Quantitative State Speciation and Extinction) was used to fit different likelihood functions of speciation on the tree for different temperature-related variables, the only significant (ΔAIC > 2) variables were: 1) mean diurnal range, and 2) maximum temperature in the warmest month (Dataset S1). We found a negative sigmoidal relationship with drift (Dataset S2) between speciation and these two variables. This means that speciation rates are higher in habitats with low diurnal temperature ranges and with mild summer temperatures, but there is a steep decline in speciation rates when the diurnal range exceeds 11°C and, separately, with summer temperatures are higher than 31°C.

PGLS and QuaSSE both found a negative relationship between speciation and mean diurnal temperature range (Dataset S1). The PGLS analysis of maximum temperature of the warmest month was not significant, but QuaSSE suggests a negative correlation as the best fit for this variable, indicating that extremely high temperatures negatively affect speciation rates. The direction of correlations between speciation and minimum temperature of the coldest month and temperature annual range (which was one of the variables with the highest *λ* for phylogenetic signal) differed between the two methods. However, PGLS P-values were significant while the QuaSSE ΔAIC was less than 2. These results suggest that milder temperatures in the coldest month and less variable temperature throughout the year are correlated with higher speciation.

The discrepancies between PGLS and QuaSSE are likely an algorithmic artifact. The PGLS used tip rate estimations extracted from BAMM, while QuaSSE evaluated correlations using its own lambda and mu estimates. Additionally, QuaSSE fits the data to five different models while PGLS is limited to a linear regression, which may not be the best model for the data.

The best fit time-dependent and (paleo)temperature-dependent model explaining Pieridae diversification dynamics was a positive exponential relationship for both speciation and extinction rates and (paleo)temperature (BEnv.VarDEnv.Var_EXPO) (Dataset S3). Global temperature, in general, underwent a negative trend during the ∼40 my of Pieridae crown group evolution. Thus, the exponential positive relationships of both speciation and extinction rates with paleoclimate means these rates also decreased (slowed) with time towards the present. The best model fit for the Pierinae tree describes a pattern in which speciation rate was constant whereas extinction rate decreased with paleotemperature (BCSTDEnv.Var_EXPO), meaning that net diversification remained positive in the group.

### Comparative phylogenetics

An ancestral state reconstruction (ASR) identified Fabales as the ancestral host plant order of Pieridae as well as the subfamilies Dismorphiinae and Coliadinae (Fig. S6). Well supported (posterior probability [pp] > 0.75) host plant switches were found in the ancestors of Pseudopontiinae (to Santalales) and Pierinae (to Brassicales). A host switch to Santalales was also detected in a clade within Pierinae that includes species-rich groups including *Mylothris, Delias*, and *Catasticta*.

*Pagel’s λ* scores suggest significant phylogenetic signal from all WorldClim climatic variables tested: mean diurnal range, maximum temperature in the warmest month, minimum temperature in the coldest month, temperature annual range and, especially, for annual mean temperature (Dataset S4). Conversely, the *Blomberg’s K* analyses were only significant for annual mean temperature and highest temperature in the warmest month, both with low P-values (<0.06) (Dataset S4).

An ASR of annual mean temperature suggests that the ancestor of Pieridae was likely found in warm climates; pronounced switches to colder, temperate climates occurred multiple times, such as in the genera *Colias* and *Pieris* (Fig. S7). The reconstruction of minimum temperature in the coldest month shows prominent habitat shifts to colder temperatures in *Colias* (Fig. S8). There was a clear shift towards habitats with greater annual temperature ranges (seasonality) in *Aporia, Colias*, and *Pieris* (Fig. S9). *Delias*, the most species-rich genus of butterflies, is often associated with warmer habitats, but many species are found in environments with cooler temperatures (Fig. S7) including tropical and subtropical montane habitats. Reconstructions of mean diurnal range (Fig. S10) and maximum temperature in the warmest month (Fig. S11) did not show pronounced variation across the tree.

### Pieridae phylogeny

We inferred the most taxonomically comprehensive phylogeny of Pieridae to date (Fig. 3, Fig. S12). We included genetic data for 593 pierid species and 15 outgroups, which amounted to a six-fold increase in taxon sampling compared to the most recent phylogeny of the family (36). Eighty-four of the 86 genera recognized in Pieridae by Lamas (31) are represented; the two missing genera, *Calopieris* and *Piercolias* are rarely collected, and the validity of the latter has been recently questioned (40). Subfamilial relationships were strongly supported (UFBoot and aLRT >95%) in general, but support for the position of Pseudopontiinae, which we recovered as the sister group to Pierinae, was low (UFBoot =75.2, SHaLRT = 77). This corroborates the results of Kawahara et al. (24) and Espeland et al. (41), based on 391 loci, but not of Chazot et al. (42), which used 9 loci (Fig. 1). Recently, Zhang et al. (40) used phylogenomic analyses to validate previous studies indicating the non-monophyly of some taxa. They suggested that *Catasticta* should be a junior synonym of *Archonias*, and that *Eurema* is paraphyletic, among other issues. In regards to *Catasticta* and *Archonias*, for example, phylogenetic, morphological, and natural history traits had already suggested that these genera are closely related to each other (38, 43, 44). However, sampling in Zhang et al. (40) was limited in some clades, and our study highlights additional issues. Namely, *Euchloe* is paraphyletic, *Rhabdodryas* is nested within *Phoebis, Glennia* is nested in *Ganyra*, and *Eurema* is still polyphyletic even after taxonomic changes by Zhang et al. (40).

**Fig. 3.**
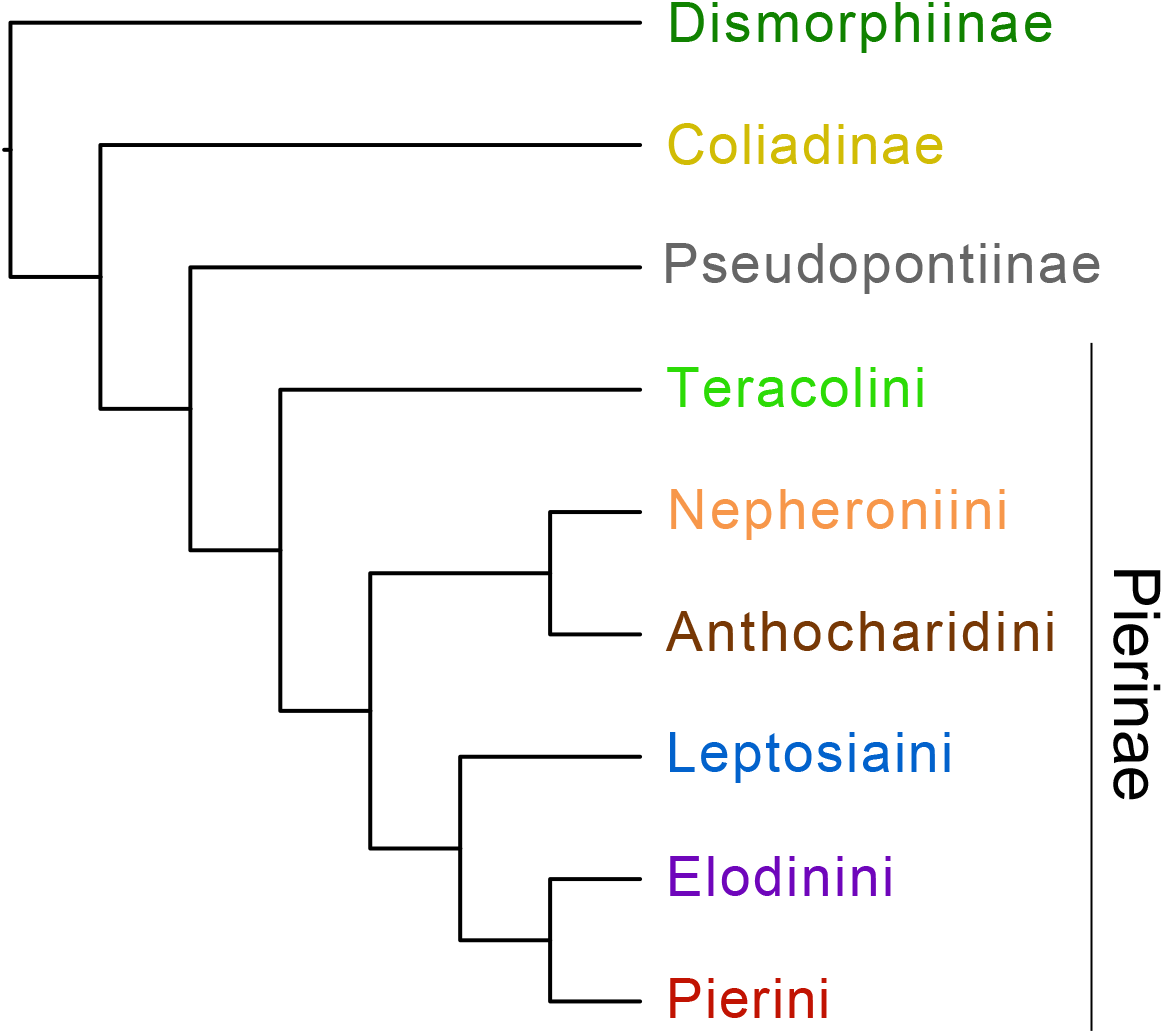
Simplified topology showing the evolutionary relationships of Pieridae subfamilies and tribes as inferred in the present study.

Divergence time estimates varied depending on the calibration scheme, with calibrations taken from Kawahara et al. (45) resulting in younger dates (Dataset S5). The crown age of Pieridae was between 69.6 – 64.5 Ma with a stem age between 83.5 – 76.1 Ma. Subfamily crown ages were estimated to be: 35.0 – 32.7 Ma for Dismorphiinae; 49.8 – 44.2 Ma for Coliadinae; and 49.9 – 48.9 Ma for Pierinae (Dataset S5). The five Afrotropical Pseudopontiinae species are on a long branch with extant taxa diversifying only 5.1 – 3.8 Ma (Dataset S5).

## Discussion

### Diversity patterns and temperature

Based on multiple analyses, our study aimed to isolate different aspects of ambient temperature and test how these could individually be correlated with rates of speciation and extinction. Our results support the hypothesis of climatic stability only when considering the range of temperature throughout the day, but not through the year (seasonality). We found evidence that contradicts the evolutionary rate hypothesis since speciation rates are lower for species inhabiting areas with high summer temperatures.

Seasonality influences species’ distributions (46). In general, diversity is greater in the tropics, and these areas are often characterized by high temperatures that are generally more constant throughout the year (47). In warm seasons, sunlight and ambient heat are the main factors influencing butterfly activity, and if conditions are not ideal, butterflies will shift the timing and length of their activity (48). The diversification of white and yellow butterflies is higher in these environments with stable circadian temperatures. However, based on our analyses, when considering seasonality (as a function of temperature annual range), we found results which do not support the hypothesis of climatic stability (12).

Warmer environments are linked to life history traits that accelerate speciation (9–11); however, our results provide evidence that in temperate environments, speciation rates are elevated. Butterflies have an optimal internal temperature range for survival, but can tolerate greater environmental ranges because they have some ability to control their thermal physiology (28, 49, 50). Nonetheless, anything below or above the tolerable range can be lethal (28). The optimal temperature range usually trends towards warmer temperatures, and most species are active during warm seasons. In some cases, the warmest month is the only time of the year when butterflies can reproduce. This is especially true for Arctic species. However, extremely warm temperatures can reduce activity in some species (48). Recognizing such trends is crucial considering global temperature changes due to climate change, particularly in light of our evidence of speciation rate decreasing with increasing temperature.

Franzén et al. (48) found that the survival of the palearctic papilionid *Parnassius apollo* is negatively affected by increased median temperatures and that there is a tendency for stabilizing selection with temperature in the lycaenid *Phengaris arion* (48). In general, if temperatures are extreme, behavior and morphology/wing coloration will adapt to regulate body temperature (51–54). Butterflies use different strategies and mechanisms to maintain body temperature, which can disrupt their regular behavior (50). They may, for example, find a shaded area to hide (28), which then prevents them from feeding and finding mates.

We found that paleotemperatures partially explain current pierid diversification patterns. Contemporary temperatures and species distributions provide only a snapshot of how temperature impacts diversity and have limited ability to explain how diversity was shaped throughout millennia. Condamine et al. (18) found that (paleo)temperature-dependent models explain diversity across a wide breadth of taxonomic groups. We found that global cooling during the Cenozoic correlates with decreases in both speciation and extinction rates across all of Pieridae. This co-variance is likely a result of high rate heterogeneity across the tree as made evident by lineage specific birth-death analyses conducted herein. However, our investigations of diversification dynamics of the most species-rich subfamily, Pierinae, show that net diversification has increased since the Cenozoic, as a result of decreasing extinction rates combined with constant speciation. Future research could further clarify these dynamics by examining smaller clades within Pieridae but with complete or nearly complete species-level sampling.

In many taxa, temperature at least indirectly correlates with diversity (6, 8, 18, 55–57). Condamine et al. (25) found evidence to support the diversification rate hypothesis in swallowtail butterflies (Papilionidae) by comparing tropical and temperate species. However, these authors caution against concluding that one trait could be responsible for explaining diversity because it could be confounded with others. In butterflies, host plant use has been frequently suggested to play an important role in diversification through coevolution (21). Wheat and colleagues found evidence in Pieridae for the signature of escalating arms races via the coupled evolution of plant chemical defenses (glucosinolates) and detoxification mechanisms (nitrite-specifier proteins) (22), as well as other key innovations involving chemical defenses derived from gene and genome duplications (23). A genus-level phylogenetic investigation of all butterflies by Kawahara et al. (24) found limited evidence that host plant switches affect diversification rates. Our exploratory ancestral state reconstruction of plant orders did not uncover any striking host shifts that coincided with diversification rate shifts, but we did not conduct any direct analyses to test for effects of host plants on diversification. Further work will be necessary to evaluate the prevalence of coevolutionary arms races along the plant/ herbivore interface in generating organic diversity.

### Comparative phylogenetics

We used *Pagel’s λ* and *Blomberg’s K* to evaluate the phylogenetic correlation of each WorldClim variable. The discrepancies between *λ* and *K* highlight the importance of assessing differences between these two methods. *Pagel’s λ* indicated phylogenetic signal consistent with a Brownian motion model for all variables. This was not the case for *K* (Dataset S4), which is a ratio of variation within clades compared to overall tree trait variation. For some of the traits that we analyzed, such as maximum temperature of the warmest month (Fig. S11), variation throughout the tree is low, but the presence of closely related taxa with very different trait values, especially on short branches (*e*.*g*., in *Pontia* + *Reliquia*) pulls *K* values down to near zero. Thus, for our dataset, *Blomberg’s K* gives the impression that phylogenetic signal is low, despite trait values being largely conserved across the tree (Fig. S11).

The general trend we observe in the Pieridae phylogeny is a uniform distribution of taxa in warm regions with mild winters (Fig. S7, Fig. S8), which is expected considering Pieridae are most diverse in warm climates. However, there are multiple independent origins of cold-adapted species (*e*.*g*., *Baltia* spp. and *Reliquia santamarta*) and clades that are sister to taxa living in warm or mild climates (such as the clade that includes *Tatochila, Hyposchila, Theochila, Pierphulia, Phulia*, and *Infraphulia*). Some of these shifts seem to be associated with adaptations to habitats at higher elevations, as shown by the convergence of morphology among groups that inhabit similarly cold, high-elevation habitats around the world (35). The monotypic genus *Reliquia*, endemic to a single mountain range in tropical Colombia, was recently found, as well as in our study, to be nested in the mostly Holarctic *Pontia* (including *Baltia*) (40). *Pontia* inhabits colder environs at similarly high elevations in the Northern Hemisphere. This observation partially uncovers the intriguing interplay of Pieridae biogeography and adaptation to high elevations around the world. Interestingly, the high-altitude, narrowly endemic monotypic Riodinidae genera *Corrachia* and *Styx*, occurring only in mountains in Costa Rica and Peru, respectively, are also nested within an Old World clade (58).

### Phylogenetic relationships of Pieridae

This study provides a robust phylogenetic framework of pierid butterflies and a foundation for testing many macroevolutionary questions. We show, with strong support (UFBoot and SH-aLRT = 100), that Dismorphiinae are sister to a clade containing Coliadinae, Pseudopontiinae, and Pierinae, in agreement with Wahlberg et al. (36) and Kawahara et al. (24). All tribal relationships of Pierinae were strongly supported and differ from previous studies (24, 36, 42). All of the Pieridae sequence data used by Kawahara et al. (24) are incorporated into our dataset, but the inferred trees differ in tribal topology (Fig. 3, Fig. S12). This is likely a result of our considerably increased taxon sampling.

We recovered the subfamily Pseudopontiinae on an extremely long branch (as shown previously in Mitter et al. (59)) of approximately 50 my in length (Dataset S5), which likely contributed to the difficulty of placing the subfamily, as well as inferring ancestral states and diversification patterns in this group. This morphologically unique Afrotropical subfamily was thought to only include a single species until 2011 (59) when molecular data demonstrated the existence of four additional cryptic species. Even though we incorporated additional molecular data for several species in the group, support for its position in relation to the other subfamilies is still low (as in Espeland et al. (41) and Chazot et al. (42)). Furthermore, several genera, like *Euchloe* and *Ganyra*, are not monophyletic (Fig. 2), and will require further taxonomic investigations.

The crown age of Pieridae of around 70 Ma overlaps the K-Pg (Cretaceous–Paleogene) boundary, a major extinction event. The lineage giving rise to the family pre-dated this event, and pierids started diversifying during the Paleogene (Fig. S1).

### Conclusions

Our study provides a robust phylogenetic framework for Pieridae and will invigorate future studies on butterfly ecology, evolution, and conservation. We demonstrate that diversity patterns in pierid butterflies are linked to current and paleoclimates, and that rates of both speciation and extinction slowed as paleotemperatures decreased during the Cenozoic.

## Materials and Methods

### Molecular data

#### 1. Locus sampling

We obtained sequence data in 3 different ways. These were: 1) the anchored hybrid enrichment (AHE) BUTTERFLY2.0 kit (60), which targets 13 genes, 2) AHE BUTTERFLY1.0 (41), which targets 425 genes, and 3) publicly available sequence data. Details of these approaches are outlined below.

##### 1.a Anchored Hybrid Enrichment

We sequenced DNA from 338 Pieridae species and two outgroups sourced from various alcoholic and dry insect collections (Dataset S6) using methods described in Kawahara et al. (24). The BUTTERFLY2.0 AHE probe set captured up to 13 loci commonly used in Sanger sequencing studies of past decades (60, 61). Raw sequencing reads underwent filtering and quality checks for base call quality (phred > 20) and length (> 30 bases) using Trim Galore! v.0.40 (62); and assembled via a custom Python v.2.7.6 script that performs an iterative baited assembly (IBA) (63) for each locus. IBA uses similarity searching in usearch (64) to assemble loci beginning with a database of reference loci from a closely related species (in this case *Danaus plexippus*) using bridger (65). Second, assembled loci were filtered by blasting the original probe regions of the assembly against the *Danaus plexippus* reference genome (66) followed by mapping single hits to check for orthology. We screened the orthologous loci for sequence contamination and removed any sequences with 99% similarity to species in another family. The resulting sequences were aligned with MAFFT v.7.0.1 (67). Different copies of DNA were collapsed into consensus sequences using FASconCAT-G v.1.0.4 (68).

We augmented the dataset with 103 pierid species sequenced for Kawahara et al. (24) with the BUTTERFLY1.0 (Dataset S6) probe set (41) which have up to 425 AHE loci (including the same 13 loci in BUTTERFLY2.0).

##### 1.b Published sequences

We complemented the AHE molecular dataset with sequences from 134 pierid species in GenBank downloaded using the software GeneDumper v.0.8 (https://github.com/sunray1/genedumper) (Dataset S6). We used the 13 loci captured by BUTTERFLY2.0 from *Colias erate* as a reference sequence for BLAST to find other available Pieridae sequences in GenBank. GeneDumper can integrate taxonomic name updates in the GenBank search and thus requires a synonym list to determine name validity. We used the species list of Lamas (31). After the original BLAST, GeneDumper filters and checks the sequence hits. When multiple sequences for the same locus of a species are available, GeneDumper considers the characteristics of the data and selects the longest sequence with the most unambiguous DNA content. The identification of the selected sequences are verified via a self-blast against GenBank’s nucleotide collection. A tiling approach is also used, where sequences are tiled along the bait sequence used in the original BLAST and the least number of tiles providing the highest coverage across the baits are chosen. Therefore, one species may be represented by sequences from multiple specimens if there are no sequences available that fully cover the bait sequence (Further details of the method can be found at https://github.com/sunray1/genedumper).

The sequences of Warren-Gash et al. (39) were made available on GenBank after we used GeneDumper, so we downloaded those sequences from the database for species not already in our molecular dataset. We also incorporated sequences from other targeted studies (38, 41), and used genomes and transcriptomes assembled to the BUTTERFLY1.0 probe kit in Kawahara et al. (24) (including two as outgroups) (Dataset S6).

There were 15 species for which we created chimeras by adding cytochrome oxidase subunit 1 (COI) from one sample to another with more loci available but no COI (see Dataset S6 for more details).

##### 1.c Combining molecular datasets

All genetic sequences were then combined to create the molecular data matrix. We undertook this process by first combining data from the two AHE datasets. The 13 loci of BUTTERFLY2.0 are included (as loci numbers 1 through 13) in BUTTERFLY1.0 (41) and simplify concatenation and combination of datasets. We then manually added GenBank sequences to this genetic dataset, and subsequently aligned each locus with MAFFT following the gene alignment protocol of Breinholt et al. (63). After alignment, each locus was examined in AliView v.1.27 (69) to check for indels, and determine reading frames. Loci were concatenated with FASconCAT-G v.1.0.4 (68) to create a concatenated dataset of 425 loci.

The final dataset had fifteen outgroups (Dataset S6) with at least two species from each of the six other butterfly families. The taxonomy of butterfly names follows Lamas (31).

#### 2. Partitioning, tree inference, and branch support

The best nucleotide partition schemes were identified using ModelFinder as implemented in IQ-TREE v.2.0.3 (70, 71). For this analysis, we allowed partitions to be identified and merged with ‘-m TESTNEWMERGEONLY’ to limit overparameterization. In order to reduce computation times, we used the rcluster method (set to 10, with a maximum of 1000) and limited the model set to GTR. We consolidated the original 425 locus partitions into 69 partitions. Separately, we ran ModelFinder to identify the best models of nucleotide evolution for our partitions using all available models. With these 69 partitions and models, we then performed 100 independent IQ-TREE tree searches with 1000 Ultrafast bootstraps (UFBoot) and 1000 Shimodaira-Hasegawa approximate likelihood ratio test (SH-aLRT) replicates to test for branch support. We implemented the ‘-bnni’ command when running these analyses to alleviate concerns about model violation inherent in the Ultrafast bootstrap method (72). All phylogenetic analyses were run on the University of Florida HiPerGator2 Cluster (www.rc.ufl.edu/services/hipergator).

#### 3. Divergence time estimation

We used two different secondary calibration schemes obtained from the trees of Espeland et al. (41) and Kawahara et al. (45) to estimate pierid divergence times (Dataset S7). Ages, based on 95% confidence intervals in these studies, were treated as hard minimum/maximum ages and we employed the penalized likelihood tree dating with TreePL (73). TreePL requires an input tree, for which we used the most likely tree (highest likelihood of 100 runs) inferred with IQ-TREE. TreePL outputs divergence times as single dates per node. In order to provide a range of dates for each divergence event, we performed TreePL independently 100 times and summarized results, following the approach of St Laurent et al. (74). This method uses several custom Python scripts (available at https://github.com/sunray1/treepl) and creates 100 bootstrap replicates generated in RAxML (75) using the 425-locus data matrix, partitioned according to the same 69 partitions inferred with ModelFinder. These 100 bootstrap alignments were then used to infer branch lengths on the constrained input phylogeny. We generated 100 bootstrap trees, utilizing optimization parameters that were determined with the ‘prime’ command and random subsample and replicate cross-validation with the ‘randomcv’ command for identifying the best smoothing parameter. This optimization and smoothing step was performed three times on each tree to identify the best combination of parameters and assess consistency in the smoothing parameter. All analyses used the ‘thorough’ command in TreePL. These analyses resulted in 100 dated trees varying only in branch lengths, which were summarized as a maximum clade credibility (MCC) tree in TreeAnnotator in the BEAST package (76). The MCC tree was used in all downstream analyses that required a dated phylogeny.

In addition to making use of secondary calibrations in TreePL, we conducted an analysis using fossil calibrations as the minimum ages. To calibrate the root, a median age of Angiosperms (139.4 Ma) from Magallón et al. (77) was used (as done by Espeland et al. (41)). In total, eight fossil calibrations were applied to the following places in the tree (fossil names and their age noted in parentheses): 1) the Parnassiinae stem age (*Thaites ruminiana*, 23 Ma); 2) the Hesperiidae stem (*Protocoeliades kristenseni*, 54 Ma); 3) an internal Hesperiidae crown age (*Pamphilites abdita*, 23 Ma); 4) Nymphidiini (Riodinidae) stem (*Theope* sp., 15 Ma); 5) the crown node of Satyrinae + Heliconiinae (Nymphalidae) (*Vanessa amerindica*, 33.7 Ma); 6) *Aporia* (Pieridae) crown age (*Aporia* cf. *crataegi*, 2.6 Ma); 7) *Belenois* (Pieridae) crown age (*Belenois crawshayi*, 0.02 Ma); 8) split between Coliadinae and (Pseudopontiinae + Pierinae) (*Vanessa pluto*, 16 Ma). Each of these fossils were selected because they have been vetted as reasonable fossils to include in calibration analyses (78). TreePL was run on this dataset as noted above, with the exception that the root was a hard maximum age and all fossils were hard minimum ages.

For the purpose of our discussion, we focus on estimated dates obtained using calibrations from Kawahara et al. (45), because this is the most comprehensive, fossil-calibrated dated phylogeny of Lepidoptera to date.

### Distribution and bioclimatic data

We used the R library occCite v.0.4.9 (79) to assemble distribution data. It provides a platform to compile georeferenced data from global databases while also generating a list of primary data sources for each record from institutional collections. We extracted occurrence data using the command ‘occQuery’ and followed the taxonomy of Pieridae according to Lamas (31). This command checks species names against the Global Names Index (gni.globalnames.org), then uses rgbif (80) to send a query to GBIF (81), requesting all records with geographic coordinates. We removed records that did not have decimal places to increase accuracy and removed records with identical coordinates to the first four decimal places because they were too precise to provide meaningful additional information. Original occurrence data sources with digital object identifiers (DOIs) were gathered using ‘occCitation’ (Dataset S8). We also extracted additional occurrence data from the literature and directly from GBIF for 37 species that were not obtained using the method described above (references also available in Dataset S8).

We mapped all records by species in an atlas using the package ggplot2 v.3.3.5 (82), with a custom R script (https://github.com/hannahlowens/PieridaeDiversity). The atlas was inspected for occurrence points that appeared as outliers or errors based on published distribution maps and opinions of Pieridae experts (authors and collaborators - see Acknowledgements) and we subsequently removed these putatively erroneous records. We also combined the records of the cryptic species *Leptidea sinapis, L. juvenica* and *L. reali* as the “sinapis complex” due to difficulty in delimiting distribution in these species (83). The final dataset had over 800,000 records for 541 pierid species, representing 91% of species in the phylogeny (Dataset S9).

We extracted bioclimatic data for the geographic locations of these records from the WorldClim dataset (2.5 arc-minute resolution; (84)) using the package raster v.3.4-10 (85). Since climatic stability can refer to different timescales including paleoclimates (86), we henceforth refer to temperature stability in regards to seasonality (annual variation) and daily range. Therefore, considering our hypotheses regarding the roles of extreme conditions and seasonality, we focused on the following variables: annual mean temperature (average of BIO1), mean diurnal temperature range (average of BIO2), maximum temperature in the warmest month (95% quantile of BIO5), minimum temperature in the coldest month (5% quantile BIO6), and temperature annual range (average of BIO7). The averages (BIO1, BIO2, BIO7), or 5% (BIO6) or 95% (BIO5) quantiles were calculated for each species that had at least one record. The data are summarized in Dataset S9.

### Host plant data

Larval host records were compiled from over 200 sources including books, websites, existing databases, and journal articles (SI Appendix), resulting in over 13,150 unique records for 347 of the 593 species in our tree. Data were captured at the taxonomic resolution provided in the original source. Many of our sources document the same butterfly-host interactions, and such duplicate records with different citations were retained, as we regarded multiple records as an indication of reliability.

Butterfly taxonomy was standardized to Lamas (31) and plant taxonomy was standardized to The World Flora (87) using taxotools v.0.0.1 (88). To provide the clearest picture of host use evolution, we elected to analyze patterns of pierid species host use at the level of plant orders rather than plant families, genera, or species. This seemed justified because many pierid species feed on Brassicaceae and Capparaceae (both Brassicales) or Santalaceae and Loranthaceae (both Santalales), a pattern noted by Ehrlich and Raven (21).

Many extensive databases of butterfly host plant records aim to aggregate all known records without scrutinizing their veracity (89, 90). In an attempt to quality-control records from the literature, we calculated the “ordinal proportion,” which is the proportion of records for a given butterfly species found in a given plant order. We scrutinized records for which the ordinal proportion was less than 20%, usually by reviewing the original references and ascertaining whether the plant order had been observed as a host plant of that butterfly species more than once. Several unusual host association records were discarded during this vetting procedure.

Even if these records are accurate, they most likely document uncommon host taxa, which are sometimes observed during population outbreaks but do not represent the most important ecological patterns (91, 92). The dataset used for the analysis is provided in Dataset S10.

### Diversification and correlation analyses

We conducted a BAMM (Bayesian Analysis of Macroevolutionary Mixtures) analysis to infer diversification rates on the pierid phylogeny. The input topology was the TreePL tree, dated with secondary calibrations from Kawahara et al. (45), excluding outgroups. We estimated prior parameters using the R package BAMMtools v.2.1.6 (93) with the command ‘setBAMMpriors’ (lambdaInitPrior ∼ 3.33; lambdaShiftPrior ∼ 0.01; muInitPrior ∼ 3.33). We ran a reversible-jump MCMC for 100 million generations, sampling every 100,000 generations with six different expected shifts (5, 10, 15, 20, 25, and 30). We used a fraction file to account for taxon sampling bias across the tree (Dataset S11). BAMM output files were analyzed in BAMMtools with a 10% burn-in. We plotted shift posterior probabilities for all six analyses and determined that they converge around 20 shifts (Fig. S13). Thus, the result based on 20 shifts was used for diversity analyses described below.

We also conducted a lineage-specific birth-death-shift (LSBDS) analysis in RevBayes (94) with an MCMC chain of 10,000 steps; following the parameters employed in Höhna et al. (95). Rate variables were set to lognormal distributions for both mean speciation and extinction rates, the rate of rate-shift events was set to uniform, and six rate categories were set as a global parameter. LSBDS does not have an option for fine level sampling bias, and therefore a global sampling fraction of 51% of 1,159 described species of Pieridae was applied.

Considering recent criticisms of estimating diversification rates using phylogenies of extant taxa (*e*.*g*., 96), we also used the castor v.1.5.7 R package (97), to estimate pulled speciation rates (PSR) and deterministic lineage through time plots (dLTT). To ensure time slices are biased more towards the present where extant time trees are expected to be more reliable, we first used the function ‘fit_hbd_psr_on_grid’ with arguments ‘Ngrid’ set to 15, and ‘Ntrials’ to 100 in order to get density values for X and Y axes. These density values were used as arguments in the castor function ‘castor:::get_inhomogeneous_grid_1D’ to create an inhomogeneous ‘age_grid’ which could then be used as an argument in the function ‘fit_hbd_psr_on_grid’ to calculate pulled speciation rate (PSR) and dLTTs on the inhomogeneous grid.

We performed comparative phylogenetic analyses using Phytools v.0.7-90 (98) in R. To estimate the degree of phylogenetic conservatism present for each of the five WorldClim BIO climatic variables, we calculated both *Pagel’s λ* (99) and *Blomberg’s K* (100) using ‘phylosig’.

A maximum likelihood-based ancestral state reconstruction (ASR) was conducted in Phytools using the ‘contMap’ function (‘anc.ML’ model) of all temperature-related variables on the dated Pieridae tree without outgroups. We also conducted a host plant ancestral state reconstruction using stochastic character mapping in *Phytools* with the ‘make.simmap’ command. For this ASR we combined five plant orders (Picramniales, Zygophyllales, Gentianales, Sapindales, Saxifragales) that were recorded as hosts for only one species each into one state (category “OTHER”) to facilitate resolving state history. Without this addition, we were unable to resolve most ancestral feeding associations. We used a symmetrical model (‘SYM’) and 1000 simulations (nsim=1000). The presence of a state was coded as 1, and the absence as 0. When there were missing data, all tested states were coded with equal prior probability. If a species was known for feeding on more than one order, we assigned equal probability for those states and 0 for all others (Dataset S11).

To assess a possible correlation between temperature and rates of diversification, we calculated phylogenetic generalized least squares (PGLS) in the R package nlme v.3.1-149 (101). Speciation and extinction rates were extracted from the results from the BAMM analysis using the ‘getTipRates’ command in BAMMtools for all species in the phylogeny. With the command ‘gls’, we set the correlation function to Brownian and method to ‘ML’. We performed ANOVA tests on resultant PGLS models to check for significance and identify the relationship of diversification rates and variable (slopes) as positive or negative.

As another means of testing for the relationship between temperature and speciation rates, we used Quantitative State Speciation and Extinction (QuaSSE) as implemented in *diversitree* v.0.9-15 (102). We used a pruned tree including only species with available trait data (541 species). Our methods largely followed Zhang et al. (103), whereby we tested increasingly complex QuaSSE models to create likelihood functions where speciation is constant, linear, sigmoidal, or hump-shaped, with and without a drift parameter. The Akaike information criterion (AIC) was used to identify the best fit QuaSSE model for each of the BIO variables.

Climatic BIO variables used in the analysis above stem from present day temperature. To evaluate how global temperature changes through geological time affected diversity in the family as well as its most diverse subfamily Pierinae, we fitted four temperature-dependent models to our phylogeny in the R. We also considered the possibility of time influencing diversity by fitting six time-dependent models. To run these analyses we used the pipeline and custom R scripts developed by Condamine et al. (104). Specifically, time-dependent models were derived from Morlon et al. (105) and Stadler (106) and the paleoclimate models from Condamine et al. (107). The original script provided by the authors included linear and exponential models, but due to criticism directed toward linear diversification dependency models by Gamisch (108) (but also see Morlon et al. (109)), we commented out code for linear models. The time-dependent models were fitted with maximum likelihood using the ‘fit_bd’ function and the paleoclimate models used the ‘fit_env’ function, both functions available in the pipeline of Condamine et al. (104). Sampling fractions of 0.5 for Pieridae and 0.44 for Pierinae were assigned to account for missing data on the phylogenies.

## Acknowledgments

David Plotkin provided suggestions on phylogenetic dating, Caroline Storer helped molecular data assembly, and Martin Wiemers helped curate distribution data. Ana B.B. Morais and Yutaka Inayoshi provided samples. Fabien Condamine made suggestions on the time- and paleotemperature-dependent analyses. Stilianos Louca assisted with the inhomogenous time grid in castor. Liam Revell provided the code for plotting temperature bars in Figure 2. We thank the NCBS Sequencing Facility for probeset sequencing of an Indian sample. We acknowledge Research Computing at the University of Florida for providing computational resources (https://www.rc.ufl.edu).

